# Benchmarking thiolate driven photoswitching of cyanine dyes

**DOI:** 10.1101/2022.09.15.507984

**Authors:** Lucas Herdly, Peter W. Tinning, Angéline Geiser, Holly Taylor, Gwyn W. Gould, Sebastian van de Linde

## Abstract

Carbocyanines are among the best performing dyes in singlemolecule localization microscopy (SMLM), but their performance critically relies on optimized photoswitching buffers. As the chemical buffer composition controls the transitions between non-fluorescent off- and fluorescent on-states, it is ultimately linked to the maximum achievable resolution. Here, we describe a workflow for screening optimal buffer conditions for single-molecule photoswitching. This is achieved by creating an intensity gradient within the field of view using a single micro-electromechanical systems (MEMS) mirror in the excitation path of the wide-field setup. Photon budget, on-state and off-state lifetimes are studied for different concentrations of mercatoethylamine (MEA) and buffer pH values. Both of MEA concentration and buffer pH determine the amount of thiolate, which is the main requirement for carbocyanine dye photoswitching. We show that thiolate acts as a concentration bandpass filter for the maximum achievable resolution and determine a minimum thiolate concentration of ∼1 mM is necessary to facilitate SMLM measurements. We also identify a concentration bandwidth of 1-16 mM in which the photoswitching performance can be balanced between high molecular brightness and high off-time to on-time ratios. Furthermore, we monitor the performance of the popular oxygen scavenger system based on glucose and glucose oxidase over time and show simple measures to avoid acidification during prolonged measurements. Finally, the impact of buffer settings is quantitatively tested on the distribution of the glucose transporter protein 4 within the plasma membrane of adipocytes. Our work provides a general strategy for achieving optimal resolution in SMLM with relevance for the development of novel buffers and dyes.

## Introduction

Single-molecule localization microscopy (SMLM) is a powerful super-resolution technique that has become a standard tool for studying biological questions (1–3). Its key advantages, i.e., sub-20 nm resolution and quantitative imaging of biological structures, is based on the precise control of the employed photoswitches. Therefore, much effort has been made to understand the photophysical and -chemical mechanisms of photoswitching (4–7).

The use of conventional organic dyes in SMLM usually requires a chemical buffer with redox properties. The first class of dyes with which reliable and highly reversible photoswitching had been reported, were the far-red emitting carbocyanines Cy5 and Alexa Fluor 647 (AF647) (8, 9). Although several other organic dyes such as rhodamine and oxazine dyes have been utilized as photoswitches in adapted chemical buffers (10–13), carbocyanine dyes still stand out for their high molecular brightness and highly reliable photoswitching performance (12, 14). These two key features have paved the way for the success of stochastic optical reconstruction microscopy (STORM) (15) and direct STORM (dSTORM) (16).

The employed photoswitching buffer usually consists of an enzymatic oxygen scavenger system and a thiol containing reducing agent, such as β-mercaptoethanol (BME) or β-mercaptoethylamine (MEA). The buffer is typically set at moderate alkaline pH to increase the formation of thiolate (RS^*−*^), which is a major compound in the creation of metastable dark or off-states (10, 13, 17), but also plays an important role in the photostabilization of organic dyes (18). In their comprehensive study, Gidi et al. describe the thiol-based photoswitching mechanism of red emitting cyanine dyes by two competing reactions, i.e., a photostability and a photoswitching pathway (19). The starting point is the reduction of Cy5 in its triplet state by RS^*−*^ and the generation of a geminate radical pair (GRP) [Cy5^*−•*^, RS^*•*^]. Hereinafter, (i) Cy5 is either restored to its ground state through back electron transfer (18) or (ii) the formation of a thiol adduct through geminate radical combination is taking place (Cy5RS^*−*^) (19). After formation of the off-state, photoinduced or thermal thiol elimination can regenerate Cy5 into its ground state, thus closing the cycle of photoswitching. In summary, the photoswitching mechanism of carbocyanines allows for (i) high photon fluxes through thiolate mediated photostabilization and (ii) creation of long lasting non-fluorescent off-states. This finally leads to the excellent performance of carbocyanines in SMLM; the detection of bright spots on the wide-field camera guarantees high localization precision (20) and transferring the majority of dyes into their off-state allows for resolving structures with high label densities.

Recently, the importance of homogeneous illumination in SMLM and its impact on single-molecule photoswitching has been demonstrated (21, 22). We have introduced a single micro-electromechanical systems (MEMS) micromirror for tunable wide-field illumination in SMLM (22). The MEMS mirror was implemented in the excitation path of our widefield setup and used as 2D scanning device of the incoming laser beam. Due to its tunability in frequency and oscillation amplitude the MEMS mirror could either be used to generate an extremely homogeneous illumination for consistent single-molecule photoswitching over large areas, or induce intensity gradients within the field of view (FOV). The latter mode allowed us to study single-molecule photoswitching at varying irradiation intensities within a single SMLM acquisition (22).

Here, the MEMS mirror is employed to study the photoswitching of AF647 on the single-molecule level by applying a range of different buffer settings, i.e., varying concentrations of MEA and pH values. We show how both parameters can be balanced towards optimal SMLM imaging conditions and characterize the working range of the associated thiolate concentration. Furthermore, we study the performance of the enzymatic oxygen scavenger system by monitoring photoswitching kinetics over prolonged imaging periods and demonstrate how the system can be stabilized when influx of oxygen is prevented.

Finally, we apply our findings by quantitatively imaging the glucose transporter type 4 (GLUT4) in the plasma membrane of adipocytes and show how buffer settings impact data quality. Insulin increases the numbers of GLUT4 molecules on the surface of adipocytes by promoting the exocytosis of GLUT4-containing vesicles from intracellular stores to the plasma membrane (23). However, recent work has argued that the spatial distribution of GLUT4 within the plasma membrane is also regulated by insulin (24, 25). Studies using dSTORM have suggested that insulin promotes the dispersal of GLUT4 from clusters to monomers, and that this may be an important aspect of GLUT4 regulation (26, 27). There is therefore a pressing need to understand the behaviour of molecules such as GLUT4 using SMLM techniques.

Our studies underpin the versatile role of thiols in singlemolecule photoswitching, provide a guideline for optimizing SMLM imaging and support the development of novel imaging buffers and dyes.

## Results and Discussion

### Impact of thiol concentration and pH on photoswitching

Single-molecule surfaces were prepared in chambered coverslips to study the photoswitching of AF647 under dSTORM conditions (22). Each chamber was filled with differently adjusted buffers, i.e., ranging pH from 6.5 to 8.5 and MEA concentrations from 10 mM to 250 mM, but utilizing the same enzymatic oxygen scavenging system. For the entire set of measurements, illumination and acquisition parameters were kept constant. Single-molecule photoswitching was studied at different laser intensities within a single FOV. This was achieved by using a certain mode of the MEMS mirror in the excitation path of our setup, thus generating a gradient with maximum laser power in the center while attenuating towards the edges of the FOV (22). Through this, the average single-molecule fluorescence could be directly related to the excitation intensity, which holds true for excitation below the saturation limit, i.e., in the lower kW cm^*−*2^ range (28, 29). In addition, it has recently been demonstrated that a reduction of the irradiation intensity of the readout laser towards sub-kW cm^*−*2^ levels has an overall positive effect on singlemolecule photoswitching and SMLM image quality (30, 31). For each buffer condition, single-molecule time traces were analyzed according to their on- and off-time intervals, which were used to determine the average lifetime of the fluorescent on- and dark off-states, τ_on_ and τ_off_, respectively (Fig. S1). In addition, the average detected spot brightness, *N*_Det_, i.e., the number of photons detected per molecule and frame, was determined. Fig. 1 shows these metrics for three different buffer conditions. Overall, increasing pH and MEA concentration led to an increase of τ_off_ and decrease of τ_on_ and *N*_Det_. Consequently, the ratio of the lifetimes, τ_off_*/*τ_on_, which is linked to the achievable resolution (12, 14, 32, 33), followed the same trend (Fig 1). These results underpin the importance of the irradiation intensity on the photoswitching metrics, with improved values towards the centre of the FOV, where the laser intensity peaked.

**Fig. 1.**
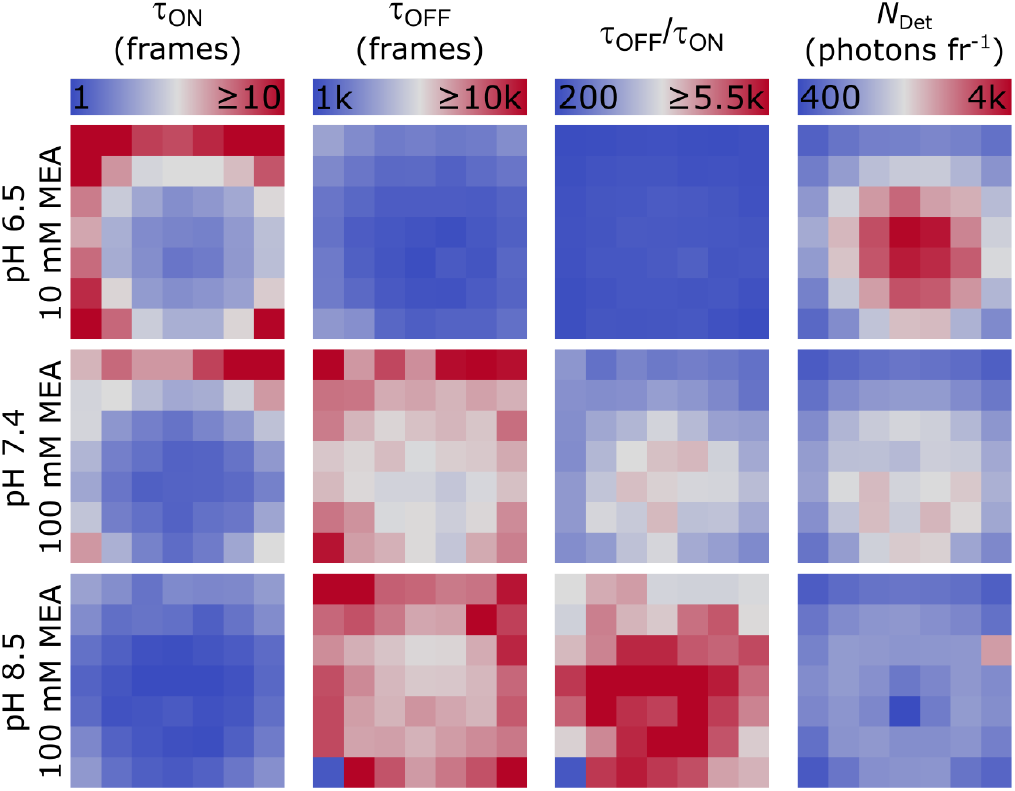
Photoswitching metrics of AF647 for different photoswitching buffer settings with sole illumination at 641 nm employing a MEMS mirror. The entire field of view (62.5 µm)^2^ was subdivided into regions of interests to account for the intensity gradient. τ_on_, τ_off_, τ_off_*/*τ_on_ and *N*_Det_ are shown for three different buffer settings, i.e., 10 mM MEA at pH 6.5 (upper panel), 100 mM MEA at pH 7.4 (central panel) and 100 mM MEA pH 8.5 (lower panel), all prepared with enzymatic oxygen scavenger system. Note the same scale for each parameter along different buffer settings. One frame corresponds to 50 ms.

Next, the rate constants at which AF647 is transferred to the offor on-state, i.e., 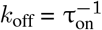 and 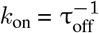, respectively, were analyzed as a function of *N*_Det_ (Fig. 2, Figs. S2,S3). Fig. 2a shows the expected linear correlation of *k*_off_ with *N*_Det_ for two different buffer conditions, i.e., 50 mM MEA at pH 6.5 and pH 7.4. The camera frame rate was chosen to sufficiently sample τ_on_ over a range of excitation intensities, hence distributing the total amount of photons emitted by the fluorophore over several consecutive camera frames. From the inverse gradient of the fit the photon budget of the fluorophore, *N*_τon_, was determined, i.e., the total number of photons emitted within τ_on_ (Fig. 2a, Fig. S2) (22). As can be seen, the gradient of the fit was decreased for pH 6.5, which corresponds to a higher *N*_τon_ when compared with pH 7.4. Next, the on-switching rate was studied for different buffer settings. Fig 2b shows the linear increase of *k*_on_ with *N*_Det_, which is linked to the photoinduced repopulation of the ground state due to thiol elimination of the dark state of AF647. Restoring fluorescence of AF647 and Cy5 has been previously demonstrated by additional excitation with blueshifted laser light, e.g., 405, 488, 514 nm, or solely by the read-out wavelength around 640 nm (14, 16, 17, 19). The latter approach is usually beneficial because of the lower sensitivity of *k*_on_ to the red-shifted excitation intensity, which allows for activating small subsets of emitters in a densely labeled sample while the majority remains nonfluorescent. Besides the photoinduced pathway, dissociation of the cyanine-thiol adduct can also occur thermally (19). Therefore, we determined the thermal recovery rate, 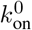, which could be extracted from the nonzero intercept of the fit as shown in Fig 2b, and finally the thermal off-state lifetime, 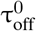. The change of intercept was already visible for a moderate pH increase, with pH 7.4 leading to a higher 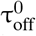 whencompared to pH 6.5. Interestingly, the gradient of the fit was significantly higher for pH 6.5, a finding which will be discussed in the next section.

**Fig. 2.**
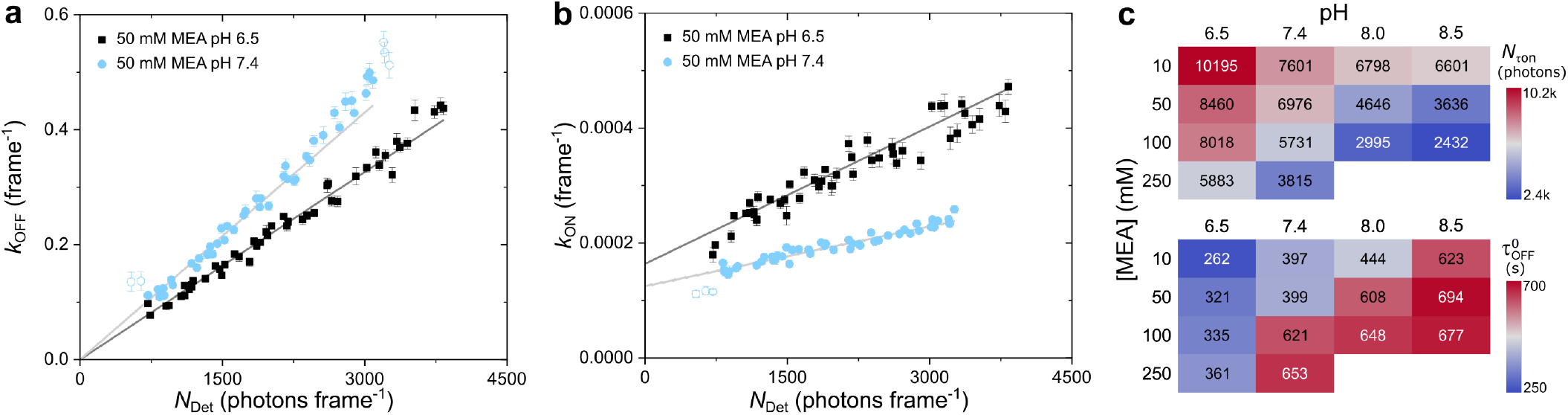
Photoswitching metrics for different buffer settings. **a**) Linear correlation of the off-switching rate, *k*_off_, and the median photon count per spot and frame (*N*_Det_, i.e., the spot brightness) for two different buffer settings, i.e., 50 mM MEA pH 6.5 (black) and 50 mM MEA pH 7.4 (blue). The photon budget of the fluorophore (*N*_τon_) was determined through the inverse gradient of the linear fit function. **b**) Linear correlation of the on-switching rate, *k*_on_, and *N*_Det_. The thermal recovery rate, 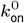, was determined from the nonzero intercept of the fit. a, b) Each data point represents a ROI as shown in Fig. 1. Linear fits to the data are shown as lines in light (pH 7.4) and dark grey (pH 6.5). Error bars are standard errors from data fits. Unfilled circles represent masked data points. One frame corresponds to 50 ms. **c**) The photon budget *N*_τon_ (top) and thermal off-state lifetime (bottom) for the entire set of MEA concentrations and pH values tested, where 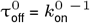.

The photon budgets and thermal off-state lifetimes for different buffer conditions are summarized in Fig. 2c. For low pH and MEA concentrations the photon budgets were higher; an increase in both parameters led to a ∼4-fold decrease from 10,200 (10 mM MEA at pH 6.5) to 2,400 photons (100 mM MEA at pH 8.5). In contrast, 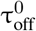 showed the opposite trend, i.e., shortened with decreasing pH and MEA concentration, e.g., from 623 (10 mM MEA at pH 8.5) to 262 s (10 mM MEA at pH 6.5). This behaviour agrees with the previous observation of proton-assisted elimination of the thiol adduct of Cy5 (19).

The ratio of τ_off_ and τ_on_ values were calculated for each ROI in the field of view and for all buffer conditions (Fig. 1, Fig. S4). τ_off_*/*τ_on_ is a meaningful parameter to determine the maximum achievable resolution with high values indicating the potential to resolve more complex structures with high label densities (14, 22, 33). This ratio is inversely linked to the duty cycle (DC = τ_on_ /(τ_on_ +τ_off_) *≈*τ_off_*/*τ_on_^*−*1^) that has also been used as a metric to characterize the photoswitching performance in SMLM (12). There are two competing effects that affect the τ_off_*/*τ_on_ ratio when varying the intensity. On one hand, an increase of the laser intensity will shorten τ_on_, which will be beneficial for the τ_off_*/*τ_on_ ratio. On the other, the sensitivity of *k*_on_ to the excitation laser as shown in Fig 2b will shorten τ_off_, which is adversely affecting τ_off_*/*τ_on_. As shown in Fig. 1 and Fig. S4a, there is a clear trend towards high τ_off_*/*τ_on_ ratios for increasing pH and MEA concentrations. τ_off_*/*τ_on_ increased through buffer adjustment by one order of magnitude from 460 (10 mM MEA at pH 6.5) to 4,600 (100 mM MEA at pH 8.5) (Fig. S4a).

### Thiolate as unique means to adjust single-molecule photoswitching

Next, we studied key photoswitching metrics as function of the thiolate (RS^*−*^) concentration, which has been identified as major compound in regulating photoswitching in organic dyes (10, 13, 19). To determine the amount of thiolate the p*K*_a_ of the thiol group of MEA needs to be known, i.e., the pH at which half of all thiols are deprotonated. Therefore, we titrated MEA in the final switching buffer, i.e., with glucose and enzymatic scavenger system (Fig. S5), and determined the p*K*_a_ of the thiol group to 8.353*±*0.004, which agrees with published values (34, 35). Using pH values of 6.5, 7.4, 8.0 and 8.5, the fractions of thiolate could then be calculated to 1.4, 10.0, 30.7, 58.4% of the applied MEA concentrations, respectively, resulting in thiolate concentrations from 0.14 to 58.4 mM as summarized in Tab. S1.

As shown in Fig. 3a the photon budget, *N*_τon_, decreased with increasing thiolate concentration, which is due to singlet state quenching and increased dark state formation (10, 19). 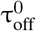 slightly increased towards ∼5 mM, then started to significantly increase and finally saturated >20 mM RS^*−*^ (Fig. 3b). The sensitivity of the recovery rate to the irradiation intensity was extracted from the gradients of the linear fits as shown in Fig. 2b. Here, a strong response of *k*_on_ could be observed for thiol concentrations <1 mM, with a 4-fold higher sensitivity at ∼0.14 mM RS^*−*^ (10 mM MEA) when compared to ∼1.38 mM RS^*−*^ (100 mM MEA), although the pH was still 6.5 (Fig. 3c). From 1 mM RS^*−*^ onwards the intensity driven response of *k*_on_ seemed to reach saturation, thus underlining the importance of a minimum concentration of thiolate to maintain long off-times during imaging at higher intensities. The τ_off_*/*τ_on_ ratio on the other hand showed an inverse dependence on the thiolate concentration when compared with *N*_τon_ (Fig. 3d). Overall, there seemed to exist no significant difference for τ_off_*/*τ_on_ and *N*_τon_, regardless whether high MEA concentrations at low pH or low MEA concentrations at high pH were used as long as the thiolate concentration remained the same, e.g., 250 mM MEA at pH 6.5 and 10 mM MEA at pH 8.0 with ∼3.5 and ∼3.1 mM RS^*−*^, respectively (Fig. 3).

**Fig. 3.**
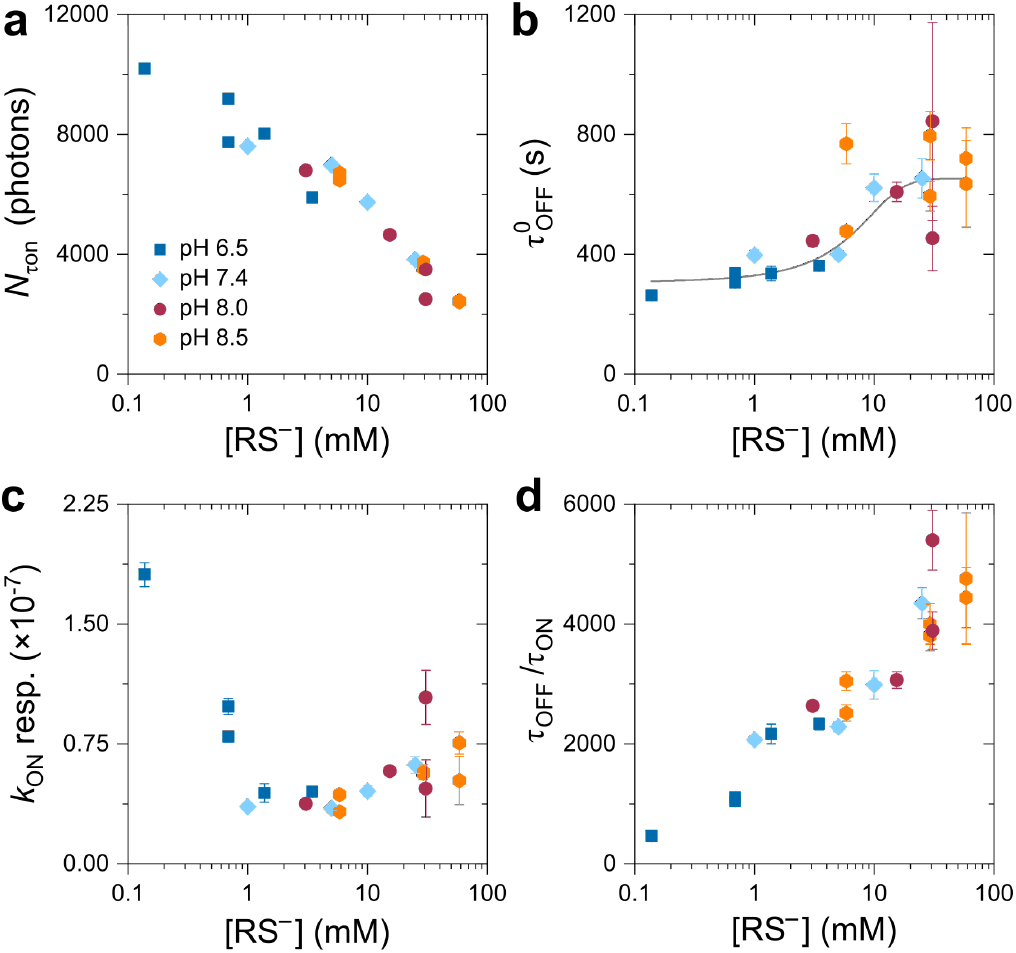
Photoswitching metrics as function of the thiolate concentration. **a**) The photon budget per on-state *N*_τon_, **b**) thermal dark state lifetime 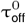, **c**) the intensity response of *k*_on_ as determined from the gradient of the linear fit as shown in Fig. 2, and **d**) the maximum τ_off_*/*τ_on_ ratio (central FOV). The different pH values are indicated by colour and symbol. The concentration of thiolate was calculated according to Eq. (1) using the experimentally determined p*K* _a_ of 8.353. Error bars are standard errors from data fits.

In SMLM and tracking experiments it is desirable to obtain high photon numbers per on-state, to allow for high localization precision (7, 36). On the other hand, the τ_off_*/*τ_on_ ratio should be as high as possible, which ensures that only single, isolated spots are localized in a densely labeled sample (14, 33). This trade off needs to be addressed by the buffer composition. Plotting both trends allowed us to identify a range for the optimal thiolate concentration (Fig. 4a) where a good photon yield and photoswitching performance can be expected, i.e., >1 mM RS^*−*^ with τ_off_*/*τ_on_ > 2,000 and *N*_τon_ > 4,000.

**Fig. 4.**
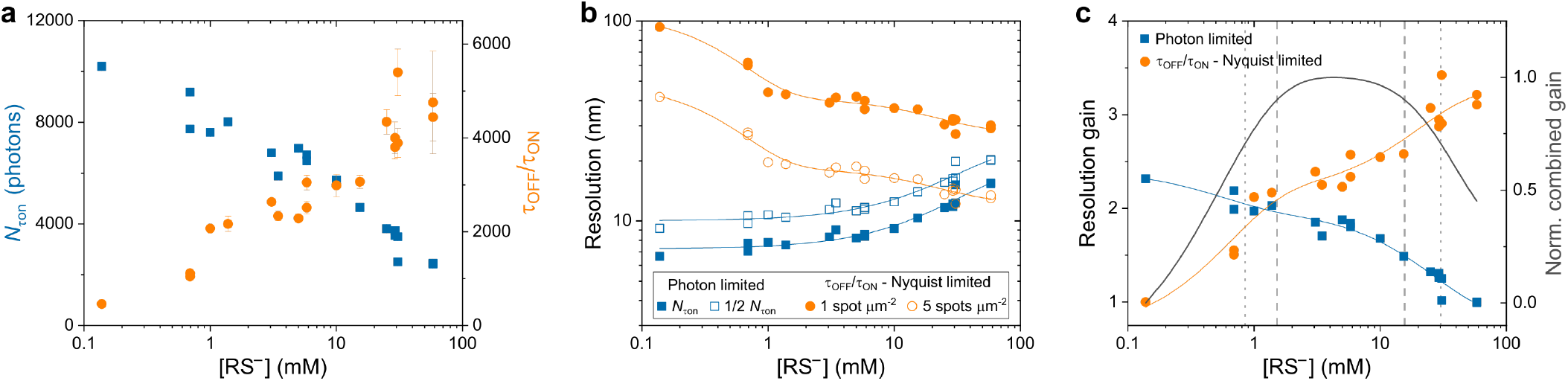
Thiolate driven achievable resolution. **a**) *N*_τon_ and τ_off_*/*τ_on_ as function of the thiolate concentration. **b**) Optical resolution calculated from the number of detected photons according to Eq. (2) (squares) and Nyquist-limited structural 2D resolution calculated from the experimental highest achievable τ_off_*/*τ_on_ according to Eq. (3) (circles). Filled squares refer to resolution determination with *N* = *N*_τon_, unfilled squares refer to 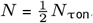. Filled orange circles refer to a label density *n* = τ_off_*/*τ_on_, unfilled circles refer to *n* = 5*×*τ_off_ */*τ_on_ ^2^ per µm^2^. Blue and orange solid lines indicate single and double exponential functions fit to data, respectively. **c**) Gain in resolution as determined by photon number (squares) and label density (circles). Solid coloured lines indicate double exponential functions fit to the data. Curve in dark grey refers to the normalized combined gain in resolution. Dotted and dashed lines in grey indicate concentration bandwidths of 0.85 and 30.15 mM (70.7% combined gain) as well as 1.5 and 15.6 mM RS^*−*^ (90%), respectively.

To make these two main parameters directly comparable, we calculated the corresponding resolution (Fig. 4b). This was done on the basis of the localization precision according to Eq. (2) and on the maximum tolerable label density based on the Nyquist criterion, Eq. (3), which states that the sampling interval must be at least twice as fine as the desired resolution (37). Here, the ability of the localization algorithm to cope with higher spot densities affects the achievable Nyquist resolution, as the localization performance can vary among different software available (38). For instance, for a label density of 2,500 µm^*−*2^ – equivalent to 40 nm 2D Nyquist-limited resolution – it makes a difference whether a localization algorithm can handle 1 or 5 spots µm^*−*2^ (Fig. 4b). In the former case τ_off_*/*τ_on_ must be at least 2,500 to ensure that one fluorophore on average will be fluorescent, whereas in the latter τ_off_*/*τ_on_ can be reduced by a factor of 5. In order to apply the Nyquist-Shannon sampling theorem to SMLM, the stochastic nature of sampling further demands an increase of the localization density (39), which underscores the importance of the τ_off_*/*τ_on_ ratio as well as the demand for robust and reliable high density localization algorithms (38). However, it has been recently demonstrated that for small inter-fluorophore distances, <10 nm, resonance energy transfers between adjacent photoswitchable dyes can lead to an increase of activation rates and thus shorten the off-times (40). For extreme cases, increasing the structural resolution through higher label densities can thus compromise the ability to resolve these.

As shown in Fig. 4b the theoretical resolution based on the available photon budget exceeded the achievable Nyquist-limited resolution. In this case the photon based resolution, which can be considered as the lower bound for the maximum achievable resolution in SMLM, remained relatively constant for thiolate concentrations <10 mM. Above this concentration this resolution significantly decreased due to thiolate induced singlet state quenching of AF647. It should be noted though that even at a thiolate concentration of 20 mM the photon number was still high enough to allow a theoretical resolution of <15 nm. This is opposite to the Nyquist resolution which increased significantly with concentrations towards 1 mM RS^*−*^, after which a further increase can be observed but at a smaller rate.

The effect of thiolate on the photon based resolution can be considered as low pass filter, excluding high thiolate concentrations as the emitter brightness is reduced. On the other hand, its effect on the τ_off_*/*τ_on_ limited Nyquist resolution can be described as long pass filter, excluding low thiolate concentrations for which high emitter densities become unresolvable. In order to evaluate the optimal thiolate concentration range, we calculated the gain in resolution from both filters (Fig. 4c). The lowest and highest thiolate concentration of the resulting passband were about ∼1 and 30 mM, respectively. Below and above this bandwidth, the benefit of one parameter is impaired by the other. Within a working concentration range of 1.5 to 15.6 mM RS^*−*^, photoswitching leads to overall high combined resolution comprising an optimal range of 2.5 and 8.3 mM (Fig. S6). The maximum gain at 4.3 mM could be realized with 43 mM MEA at pH 7.4. At this pH the fraction of thiolate is 10% of the employed MEA concentration and thus allows for tuning the thiolate concentration conveniently within the concentration bandwidth. Table 1 and Fig. S6d summarize the photoswitching performance of AF647 with MEA.

**Table 1.** Optimal buffer conditions as used in this study for the carbocyanine dye AF647. Performance was rated with + (very good), o (decent), *−* (bad). For conditions rated + the buffer contained between 1.5 and 15.6 mM RS^*−*^ (cf. Fig. S6).

| [MEA]<br>(mM) | pH |  |  |  |
| --- | --- | --- | --- | --- |
|  | 6.5 | 7.4 | 8.0 | 8.5 |
| 10 | – | o | + | + |
| 50 | – | + | + | o |
| 100 | o | + | – | – |
| 250 | + | o |  |  |

Buffers in SMLM experiments should hence be prepared to realize the major gain in structural resolution while avoiding a significant loss of localization precision. With the τ_off_*/*τ_on_ ratio as the main limitation to resolve densely labeled structures, it therefore makes sense to exceed the minimum thiolate concentration of *≤*1 mM. If the structural complexity, however, is low the thiolate concentration might be reduced to ≤1 mM to allow for high localization precision and fast data acquisition (Figs. 3, 4). On the other hand, buffers with low thiolate concentrations could be advantageously used for imaging complex structures with superresolution techniques that rely on the temporal analysis of intensity fluctuations such as in SOFI (41) or SRRF (42).

### Acidification of the enzymatic oxygen scavenger system

The oxygen scavenger system applied in this study was based on glucose oxidase and catalase (GOC), which catalyzes the reaction of D-glucose and oxygen to gluconic acid (43). It is a well-established system for single-molecule experiments in aqueous environment, which facilitates efficient depletion of oxygen within seconds (44). Accumulation of gluconic acid over time, however, results in acidification of the solvent if oxygen can re-dissolve from the headspace (45). This has motivated the development of alternative systems such as substituting glucose oxidase with pyranose oxidase (46).

Here, we investigated the photoswitching of AF647 over prolonged time periods. Because a pH drop over time would affect the photoswitching performance due to a reduction of the thiolate concentration, we imaged a single-molecule surface in steps of 2 hours after preparing the buffer (50 mM MEA, pH 7.4) and left the chamber unsealed. We measured an increase of τ_on_ within four hours from 161 (0 h) to 183 (2 h) and 224 ms (4 h), while τ_off_ decreased from 169 (0 h) to 97 (2 h) and 55 s (4 h) due to proton-assisted restoration of the ground state of AF647; the pH dropped by ∼1 every two hours (Fig. 5, left). Here, the acidification of the GOC system was slower than previously reported (46), which can be explained by the presence of MEA with its main buffering capacity around pH 8.35.

**Fig. 5.**
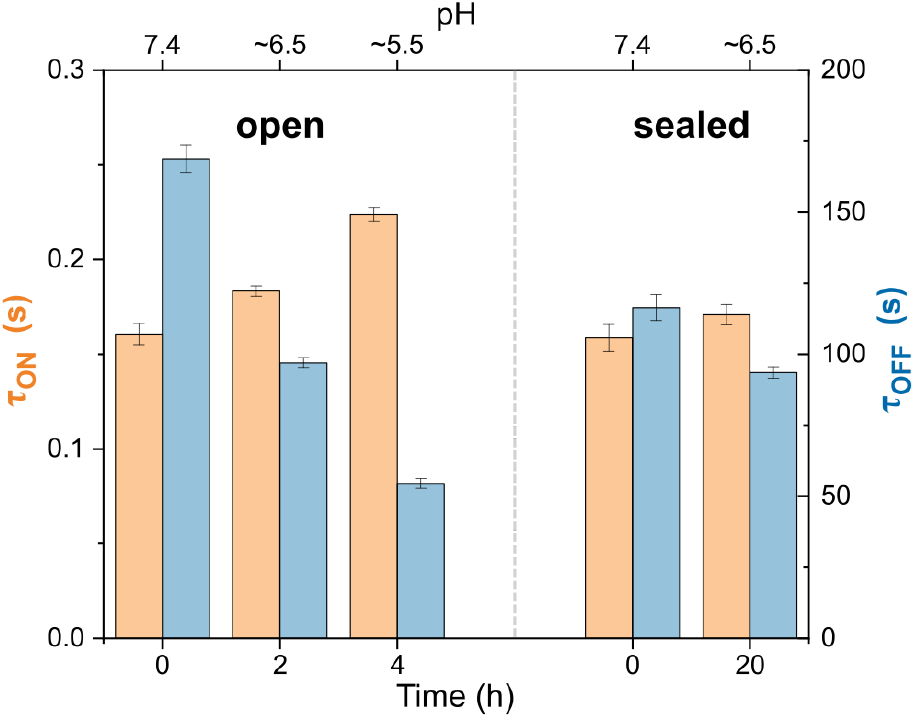
Evaluating the stability of the photoswitching buffer (50 mM MEA, pH 7.4, glucose, glucose oxidase and catalase system) in sealed and unsealed sample chambers. τ_on_ (orange) and τ_off_ (blue) were determined for each measurement. Left: unsealed LabTek chamber imaged just after adding the buffer (0 h, pH 7.4), 2 h (pH ∼6.5) and 4 h (pH ∼5.5) afterwards; right: completely filled and sealed LabTek chamber just after adding the buffer (0 h, pH 7.4) and 20 h afterwards (pH ∼6.5). Measurements were performed with flat-field illumination. Error bars are standard errors from data fits.

If the reaction chamber was completely filled with photoswitching buffer and sealed on top with a glass coverslip to avoid any headspace for gas exchange, acidification could be dramatically reduced as shown in Fig. 5 (right). τ_on_ and τ_off_ changed by 8% and -20% after an incubation time of 20 hours at room temperature, which is significantly lower compared to the unsealed chamber (39% and -68%, respectively, after 4 hours). The pH was measured to drop by ∼1 within 20 hours, thus the stability was increased by at least one order of magnitude. Our results agree with the findings employing GOC with 143 mM BME at pH 8 where single-molecule brightness and localization density were studied in chambers sealed with parafilm (30). pH stability could be further increased by increasing the buffering capacity of the thiol compound, which can be achieved by increasing the thiol concentration or setting the pH towards the p*K*_a_.

Single-molecule photoswitching has also been demonstrated using solely MEA at moderate alkaline pH in aqueous environment (10, 14, 47). Besides its photoswitching capability, it has further been demonstrated that MEA solutions also act as efficient oxygen scavenger although at depletion rates lower than the GOC system (44). We therefore used our approach to compare photoswitching of AF647 in presence and absence of GOC (Fig. S7). Both *N*_τon_ and τ_off_*/*τ_on_ are moderately improved by using GOC for the optimal thiolate concentration as determined in this work, thus its use in SMLM is generally advisable. However, the use of a buffer with mere thiol can be beneficial for applications where the photoswitching buffer should be designed as simple as possible, e.g., spectroscopic measurements (13) and correlative microscopy approaches where SMLM is combined with atomic force microscopy (47) or expansion microscopy (48).

### Consequences for biological imaging

To demonstrate the impact on minor changes of the pH in biological imaging, we imaged the glucose transporter GLUT4 in the basal membrane of adipocyte cells in flat-field illumination with 100 mM MEA (Fig. 6). To assess the overall SMLM image quality, we determined the resolution by Fourier ring correlation (FRC) (49), which was calculated to 26*±* 3 and 34*±* 5 nm (median*±*MAD) for pH 7.4 and 8.0, respectively. The decrease in FRC resolution can be explained with a reduced amount in localizations from a density of ∼1,740 to ∼560 localizations µm^*−*2^. Consequently, GLUT4 should be imaged at pH 7.4 under the experimental settings. The corresponding thiolate concentration of 10 mM is within the proposed concentration range and allows for imaging at high τ_off_*/*τ_on_ ratio, whereas an increase of the pH by 0.6 to pH 8.0 exceeds the thiolate concentration by a factor of three (∼31 mM) with an overall detrimental effect on the resolution.

**Fig. 6.**
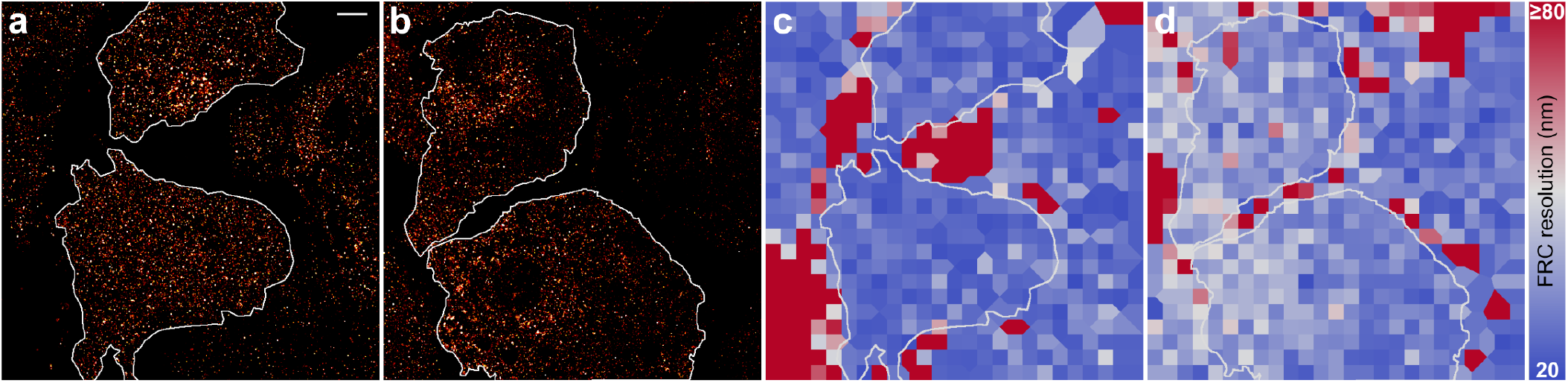
SMLM imaging of GLUT4 in the basal membrane. **a**) dSTORM image applying 100 mM MEA at pH 7.4 and **b**) 100 mM MEA at pH 8.0 employing the GOC system. Two cells for each condition were selected and analysed as shown as white marking. **c, d**) The corresponding FRC maps with the colour code set to FRC resolutions between 20 and *≥*80 nm. The FRC resolution was determined to c) 26.1 *±* 3.1 (median *±* MAD, top cell) and 26.4 *±* 2.8 nm (bottom cell), and d) 35.1 *±* 4.3 (top cell) and 32.8 *±* 5.0 nm (bottom cell). The scale bar of 5 µm in a) applies to all images.

The resolution as determined by FRC could be increased for higher pH values by increasing the acquisition length to gain more localizations. Alternatively, additional activation at shorter wavelength could be used (16, 17, 19). On the other hand, the increased τ_off_*/*τ_on_ ratio for pH 8.0 allowed for imaging with a reduced amount of artefacts, but whether the label density demands this ratio is questionable. The τ_off_*/*τ_on_ ratio always sets a limit for the maximum achievable resolution, but it should ideally fit the label density to not waste acquisition time. This is, however, technically challenging without a priori knowledge of the label density.

In summary, for achieving the highest possible the resolution the thiolate concentration should be kept as low as possible to gain high photon budgets and the maximum amount of localizations. It is advisable to adapt the thiolate concentration a posteriori, e.g., if artefacts are detected (49). To minimize artefacts from the very start, thiolate can be used at higher concentrations, but this can come at the cost of an overall reduced resolution.

## Conclusions

Controlling the photoswitching performance is key for the successful implementation of SMLM in quantitative biology. Therefore, the right adjustment of the chemical buffer composition is of utmost importance. By means of a MEMS micromirror we studied single carbocyanine dye molecules at varying intensities within a single acquisition and by using an automated workflow we determined key photoswitching metrics such as spot brightness, *N*_Det_, the fluorescent on-state lifetime, τ_on_, and off-state lifetime, τ_off_, for a range of different buffer settings. The performance of photoswitching was linked to the thiolate concentration, which is defined by the concentration of the thiol such as MEA and the pH of the solvent.

Thiolate was characterized to have a concentration bandwidth in the range of 1-16 mM, which allows for achieving high resolution through molecular brightness and τ_off_*/*τ_on_ ratio. This range allows for tailoring carbocyanine photoswitching to imaging needs. Reducing the thiolate concentration will enhance molecular brightness, whereas an increase shortens τ_on_, improves the longevity of τ_off_ and thus the duty cycle. A good starting point will be 50 mM MEA at pH 7.4 or 10 mM MEA at pH 8.0 with ∼5.0 and ∼3.1 mM thiolate, respectively. Although suboptimal for SMLM, minimal thiolate concentrations can be interesting for methods based on temporal analysis of intensity fluctuations (41, 42).

The optimal thiolate concentration for cyanine photoswitching can be realized at different pH values (Fig. 3). This will allow for setting the pH towards levels that are facilitating other chemical processes in the sample, e.g., enzymatic activity or the correlative use of dyes with pH dependent blinking (50, 51). This way, the ionic strength of the buffer can be varied while maintaining the photoswitching performance. For a comparison of different thiol containing reducing agents the corresponding thiolate concentration needs to be taken into account. Besides MEA, BME is frequently used for photoswitching of organic dyes (12, 19, 30), typically at concentrations of 1%, i.e., 143 mM at pH 8.0. With a p*K*_a_ of 9.6 for BME (52) the corresponding thiolate concentration can be calculated to ∼3.5 mM, which is comparable with the aforementioned MEA concentrations. However, the thiolate concentration can only be calculated using a precise estimate of the p*K*_a_ of the thiol group, and a range of values has been published owing to different measurement techniques and experimental conditions, e.g., for MEA 8.19 (53), 8.27 (54), 8.31 (55), 8.35 (34, 35), 8.4 (56), and 8.6 (57), and for BME 9.5 (57), 9.61 (52), 9.72 (53, 54), 9.8 (56). The small deviations among these published values can render an estimation of the exact amount of thiolate challenging, e.g., with ∼2.4 fold more thiolate at pH 7.4 when using 8.19 instead of 8.6 as p*K*_a_ (cf. Fig. S8). As such, a titration of the employed thiol in the final experimental buffer environment is advisable. This also underpins the importance of indicating pH and buffer conditions in photoswitching experiments.

Acidification of the popular oxygen scavenger system employing glucose, glucose oxidase and catalase has been investigated in several studies (30, 45, 46, 58). Here, we demonstrated that sealing of the reaction chamber without leaving air headspace is the most critical measure to avoid a significant pH drop with GOC over time. By monitoring τ_on_ and τ_off_ over time AF647 has been used as a sensor for the pH change. By proper sealing, e.g., by using a coverslip on top of a fully filled imaging chamber, consistent photoswitching can be performed for several hours without significant pH drop even with the GOC system. Alternative oxygen scavenger systems without being prone to acidification can hold the pH, but if the imaging chamber remains unsealed the chemical composition will change over time due to the ongoing influx of oxygen.

Ultimately, the perfect buffer composition will depend on the structural complexity in SMLM experiments. High τ_off_*/*τ_on_ ratios are needed when the label density increases, which can be the case for biological samples with a natural variability in protein expression. Therefore, a perfusion system could advantageously be used to adapt the amount of thiolate on the fly (59), i.e., through changing pH or thiol concentration.

## Materials and Methods

### SMLM setup

The setup was a single-molecule sensitive wide-field microscope equipped with a single MEMS micromirror, which has been previously described in detail (22). Use of a NA 1.49 60*×* oil immersion objective (APON60XOTIRF, Olympus) and a ∼1.8*×* post magnification (OptoSplit II, Cairn) led to an effective camera pixel size of 122 nm pixel. The fluorescence light was filtered with a zt532/640rpc dichroic mirror (Chroma) and multibandpass filter ZET532/640 (Chroma) and imaged on a EMCCD camera (iXon Life 888, Andor). For all measurements, the camera was recording 30,000 frames at 20 Hz frame rate, except for the experiments on buffer acidification with 15,000 frames at 10 Hz. Buffer acidification and cell measurements were performed with a refractive beam shaping device to generate a flat-field illumination (piShaper 6_6_VIS, AdlOptica), whereas all other experiment were performed with active MEMS micromirror. The excitation intensity for the full FOV was measured to 0.48 kW cm^*−*2^ for singlemolecule photoswitching using the MEMS illumination and 0.72 kW cm^*−*2^ for cell measurements using the piShaper.

### Sample preparation

For preparing single-molecule surfaces we used the following complementary DNA sequences purchased from Eurogentec Ltd: 5’-GGGAATGCGAATCAAGTAATATAATCAGGC-3’, which was biotinylated at the 5’ end, and 5’-GCCTGATTATATTACTTGATTCGCATTCCC-3’, which was modified with AF647 at position 8 via internal labelling. Hybridization to dsDNA was performed by mixing sense and antisense strand at a ratio of 2:1 and incubating overnight at room temperature. Singlemolecule surfaces were prepared on the basis of albumin, biotinylated albumin and NeutrAvidin as previously described (22).

3T3-L1 adipocytes that stably express a version of GLUT4 which includes a HA epitope in the exofacial domain that is accessible to antibodies in intact cells only when GLUT4 is at the cell surface were grown and differentiated on Nunc multichambered slides as described (27). HA-GLUT4GFP is a well-used reporter for GLUT4 trafficking (60–62). Adipocytes were incubated in serum-free media for 2 h then fixed with 4% paraformaldehyde (PFA) in PBS for 20 min at room temperature. The samples were quenched with 50 mM NH_4_Cl in PBS for 10 min, washed with PBS then incubated in blocking solution (2% BSA with 5% goat serum in PBS) for 30 min prior to incubation with anti-HA antibodies (Covance product MMS 101P, mouse monoclonal HA.11) for 90 min. After washing cells were incubated with AF647 antimouse antibodies (1:1000) for 60 min, washed and then incubated in PBS prior to analysis.

### Photoswitching buffer

The photoswitching buffer was prepared according to (44): the enzymatic oxygen scavenger system consisted of 5% (w/v) glucose, 10 U mL^*−*1^ glucose oxidase, 200 U mL^*−*1^ catalase in PBS (all from Sigma); mercaptoethylamine (MEA, purchased as cysteamine hydrochloride, Sigma) was added to the solution at final concentrations of 10–250 mM as indicated; finally the pH was adjusted by adding designated amounts of KOH, except for 10 and 50 mM at pH 6.5 where HCl was used. If not otherwise stated, the LabTek chambers (Nunc) were completely filled and sealed with a coverslip on top to avoid further gas exchange and air bubbles.

For the titration curve a 100 mM MEA solution was prepared in GOC photoswitching buffer as described above (Fig. S5). 40 mL of the freshly prepared solution was titrated at room temperature with 1 M KOH while the pH was monitored by a pH meter (Oakton pH 700); the solution was thoroughly mixed via magnetic stirrer and stir bar.

The thiolate concentration was determined according to the Henderson-Hasselbalch equation (63):

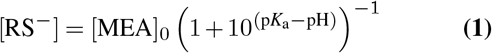

with [MEA]_0_ = [RS^*−*^] + [RSH] as the total concentration of thiol used.

For monitoring the time-dependent acidification of the photoswitching buffer, the pH of the solvent in the LabTek chamber was measured using pH indicator strips (pH range 5.0-10.0, MQuant) with a step reading of pH 0.5. Photoswitching buffer was prepared with GOC system and 50 mM MEA and set to pH 7.4 using KOH. Measurement of the pH was done at each time point. Unsealed specimen were filled with 750 µL of switching buffer and imaged with the lid off with data being taken just after adding the buffer (0 h), two and four hours later. The specimen was not moved between experiments and kept on the microscope. Sealed specimen was prepared in the same way, but the chamber was completely filled and sealed with coverglass on top. The pH was measured just before sealing and after imaging at 20 hours later.

### SMLM data analysis

The analysis of photoswitching kinetics was performed as previously described (22). Briefly, raw acquisitions were processed with rapidSTORM 3.3 (64) and subsequently analysed with Fiji (65) and custom written ImageJ macros (66). The resulting localization files were then loaded into Fiji with 10 nm pixel resolution for the reconstructed images. The image was subdivided into 7*×*7 ROIs. For the geometrical inspection of localizations originating from single-molecule photoswitching (localization pattern), masks were created through Gaussian smoothing with 1 px standard deviation. Only localization patterns were selected if mask had a minimum circularity of 0.8 and a pixel area between 3 to 300. Raw localizations within selected localization patterns were analyzed with respect to the molecule’s on- and off-time intervals. For each ROI on- and off-time histograms were generated and fit to an exponential function to obtain the characteristic lifetimes of on- and off-state, respectively (cf. Fig. S1 and (22)). For the on-time analysis, we allowed our algorithm to tolerate a gap of four frames between consecutive localizations to be tolerated, which was empirically found through an incremental increase of the gap interval until saturation (22). The photon count of each spot was extracted from the 2D Gaussian fit of the localization software, which is known to underestimate the total number of photons detected (67, 68), but for single-molecule surfaces this mismatch can be considered constant throughout the data set. The spot brightness was determined as the median photon count of all localizations passing the selection within a single ROI. Maps as shown in Fig. 1 were generated with the obtained photoswitching values for each ROI. For the kinetic analysis as shown in Figs. S2 and S3 outliers or values where τ_on_ was undersampled, which could occur in high intensity ROIs, were excluded from analysis (22). This was necessary for higher MEA concentrations and pH values. Numeric values as summarized in Fig. 2c and Fig. S4 were obtained from data fits and averaged if more than one acquisition per condition was made. Fourier ring correlation (FRC) maps were generated using the ImageJ plugin NanoJ SQUIRREL (49). The theoretical resolution was determined from the variance of the localization uncertainty 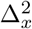.

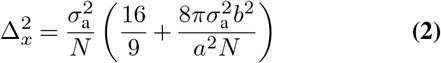

With 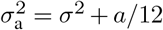 and *σ* as the standard deviation of the point spread function, *a* as the camera pixel size, *b* as background noise and *N* as number of detected photons (67). The precision was calculated as the square root of twice the variance to account for the excess noise of the EMCCD (67). The theoretical precision was multiplied with 2.355 to obtain the theoretical resolution (Fig. 4b, circles). For the data shown in Fig. 4, *σ* was set to 140 nm, *a* = 120 nm, *b*^2^ = 49 photons; for *N* two scenarios were used: *N* = *N*_τon_ and 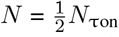. The former scenario ideally assumed that all photons would be captured within one frame leading to a single localization, whereas the latter refers to a more realistic spot brightness due to uncorrelated fluorescence emission and frame acquisition.

The two-dimensional structural resolution was calculated according to the Nyquist-Shannon sampling theorem (37).

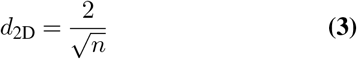

with *n* as the label density. The τ_off_*/*τ_on_ ratio was considered as the upper limit for the maximum tolerable density within the diffraction limited region (DLR). The DLR is sometimes considered as the FWHM of the PSF, e.g., 340 nm, which would in principle allow to resolve eleven spots per µm^2^. Localization software, however, fails in detection of such a high emitter density even if high density algorithms are used. We therefore used a more conservative approximation of the DLR, i.e., DLR = 1 µm^2^, which also allowed to use *n* = τ_off_*/*τ_on_ in Eq. (3) (Fig. 4b, squares). In addition, a high density case was considered with 5 spots µm^*−*2^ (38).

## ACKNOWLEDGEMENTS

We are grateful to Ralf Bauer and Paul Janin (University of Strathclyde, Glasgow) for providing us with the MEMS mirror. This work was supported by the EPSRC (EP/V048031/1), the Academy of Medical Sciences/the British Heart Foundation/the Government Department of Business, Energy and Industrial Strategy/the Wellcome Trust Springboard Award (SBF003\1163). LH was supported by a doctoral fellowship (EPSRC studentship 2031229 and EPSRC Doctoral Training Partnership (DTP) grant EP/N509760/1). HT was supported by a Studentship from Diabetes UK (18/0005905) and AG by a studentship from EPSRC (EP/T517938/1).

## Supplementary Information

**Fig. S1.**
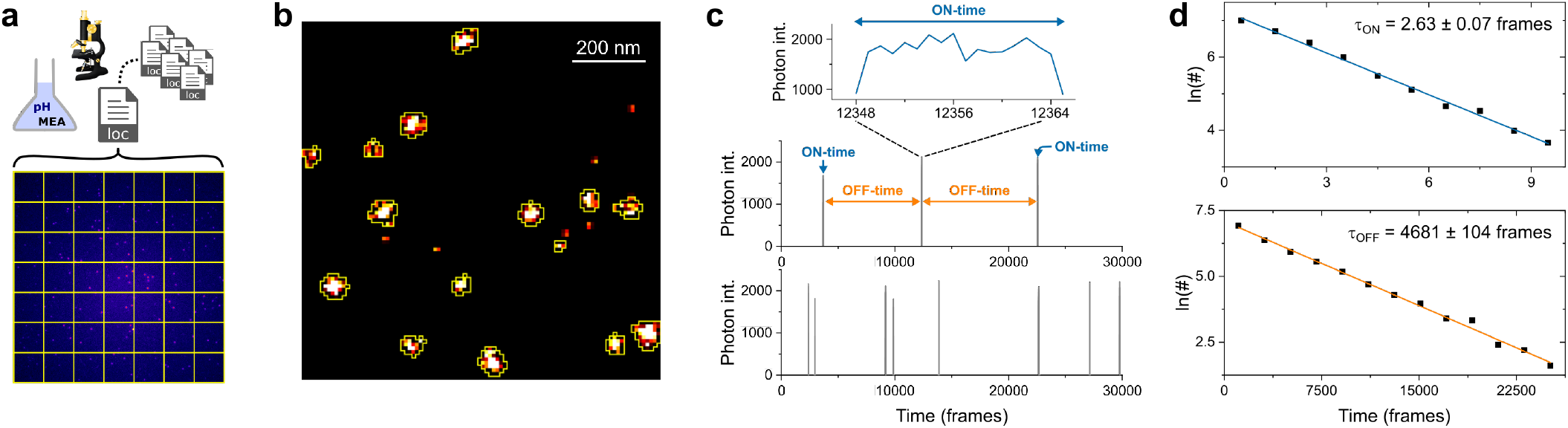
Single-molecule photoswitching analysis. **a**) For every buffer settings full FOV measurements of single-molecule surfaces were performed under dSTORM conditions (bottom). The obtained localizations were subdivided into different regions of interests (ROIs). b. Single localization patterns in the dSTORM image were subject to geometrical inspection; if successful (as indicated by the yellow selections) the corresponding localizations from the raw file were prepared as single-molecule time trace. The exemplary dSTORM image shows a small section of a single ROI. **c**) Time traces of each localization pattern were analyzed according to on- and off-time intervals, which were summed into an on-state and off-state histogram per ROI, respectively. **d**) The histograms were fitted to a single exponential decay using the function ln *y* = ln *a − kx*, with *k* as rate constant and 1*/k* as the characteristic lifetime τ. One frame corresponds to 50 ms.

**Fig. S2.**
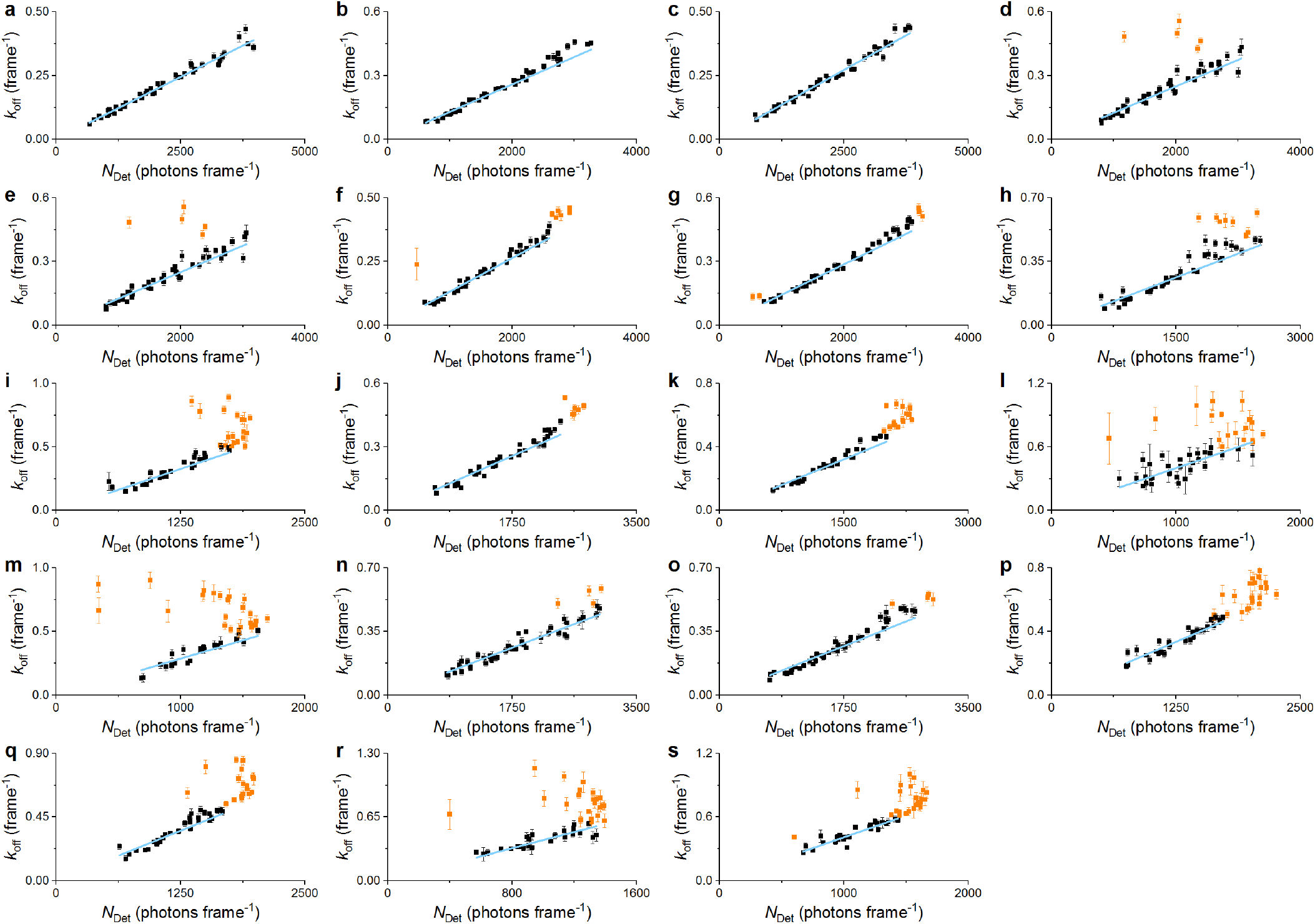
Linear correlation of *k*_off_ and *N*_Det_ for various buffer conditions. All buffer settings with enzymatic oxygen scavenger system. a) 10 mM MEA pH 6.5, **b, c**) 50 mM MEA pH 6.5, **d**) 100 mM MEA pH 6.5, **e**) 250 mM MEA pH 6.5, **f**) 10 mM MEA pH 7.4, **g**) 50 mM MEA pH 7.4, **h**) 100 mM MEA pH 7.4, **i**) 250 mM MEA pH 7.4, **j**) 10 mM MEA pH 8.0, **k**) 50 mM MEA pH 8.0, **l, m**) 100 mM MEA pH 8.0, **n, o**) 10 mM MEA pH 8.5, **p, q**) 50 mM MEA pH 8.5, **r, s**) 100 mM MEA pH 8.5. Fit function is shown as blue line. From the gradient of the fit the photon budget *N*τon was determined. Extreme outliers and values subject to undersampling, shown in orange colour, were excluded from analysis (cf. (22)). One frame corresponds to 50 ms. Error bars are standard errors from data fits.

**Fig. S3.**
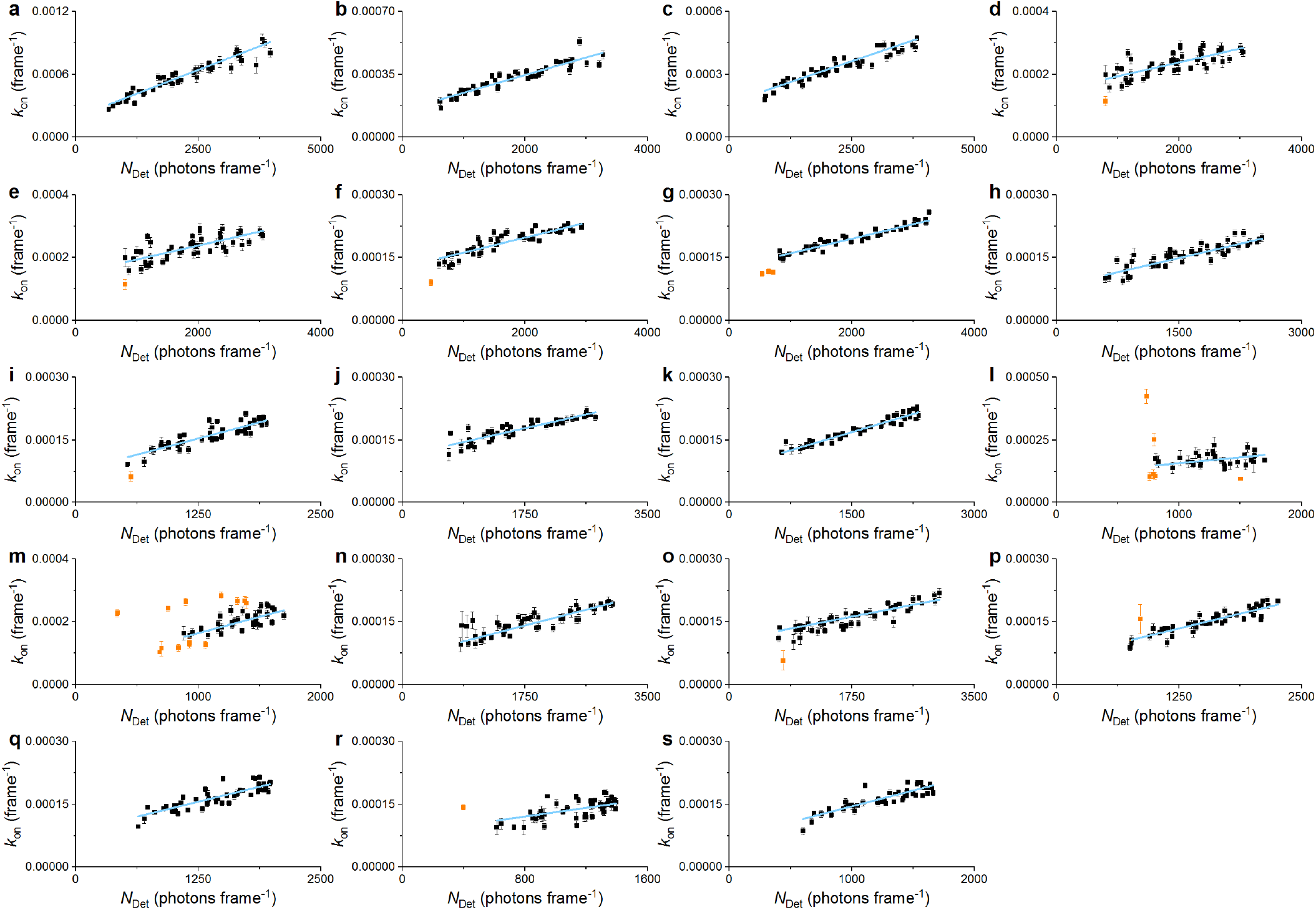
Linear correlation of *k*_on_ and *N*_Det_ for various buffer conditions. All buffer settings with enzymatic oxygen scavenger system. **a**) 10 mM MEA pH 6.5, **b, c**) 50 mM MEA pH 6.5, **d**) 100 mM MEA pH 6.5, **e**) 250 mM MEA pH 6.5, **f**) 10 mM MEA pH 7.4, **g**) 50 mM MEA pH 7.4, **h**) 100 mM MEA pH 7.4, **i**) 250 mM MEA pH 7.4, **j**) 10 mM MEA pH 8.0, **k**) 50 mM MEA pH 8.0, **l, m**) 100 mM MEA pH 8.0, **n, o**) 10 mM MEA pH 8.5, **p, q**) 50 mM MEA pH 8.5, **r, s**) 100 mM MEA pH 8.5. Fit function is shown as blue line. From the *y*-intercept the thermal dark state 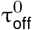 was determined. Extreme outliers and values subject to undersampling, shown in orange colour, were excluded from analysis (cf. (22)). One frame corresponds to 50 ms. Error bars are standard errors from data fits.

**Fig. S4.**
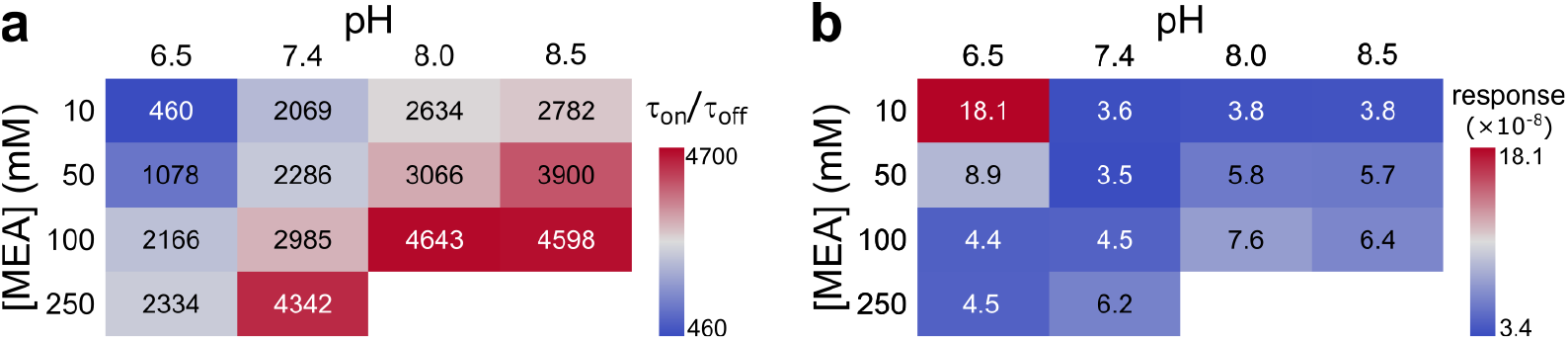
**a**) The maximum achievable τ_off_*/*τ_on_ in the FOV for different MEA concentrations and pH values. **b**) The response of *k*_on_ (frame*−*1) on the spot brightness *N*_Det_ as determined from the gradient of the linear fit in Fig. 2b.

**Fig. S5.**
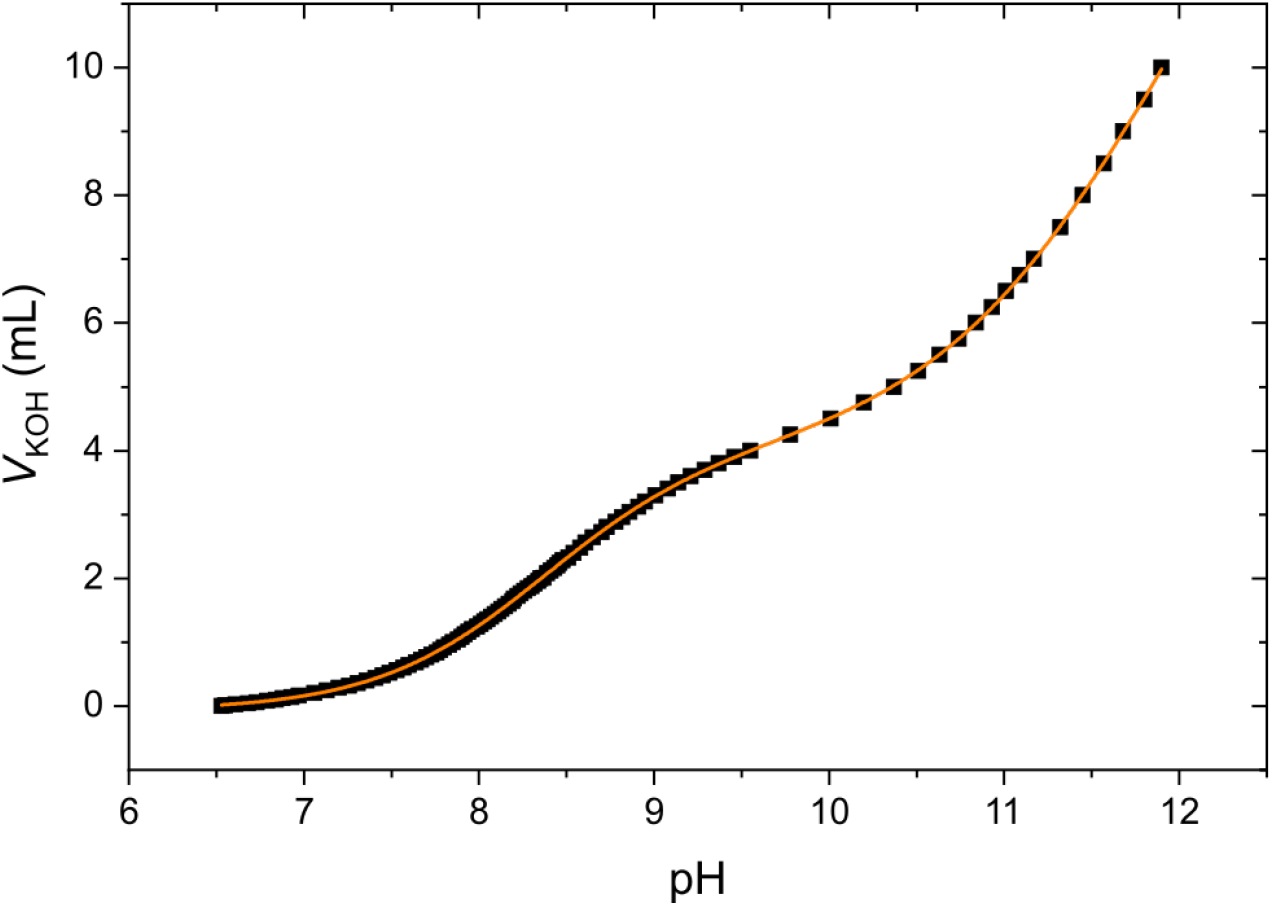
Titration of MEA. A solution of 100 mM mercaptoethylamine (MEA) hydrochloride was prepared in photoswitching buffer, i.e., 5% glucose, 10 U mL*−*1 glucose oxidase, 200 U mL*−*1 catalase, complemented with phosphate buffered saline, which was also used for the single-molecule imaging experiments in this work. Titration was carried out with 1 M potassium hydroxide (KOH). A sum of two Boltzmann functions was fit to the data to determine the p*K* a of the thiol and amino group to p*K* a1 = 8.353 *±* 0.004 and p*K* a2 11.931 *±* 0.033, respectively (R2 = 0.99998); 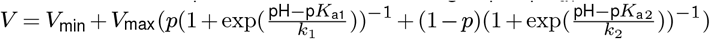. The fit curve is shown as orange line.

**Fig. S6.**
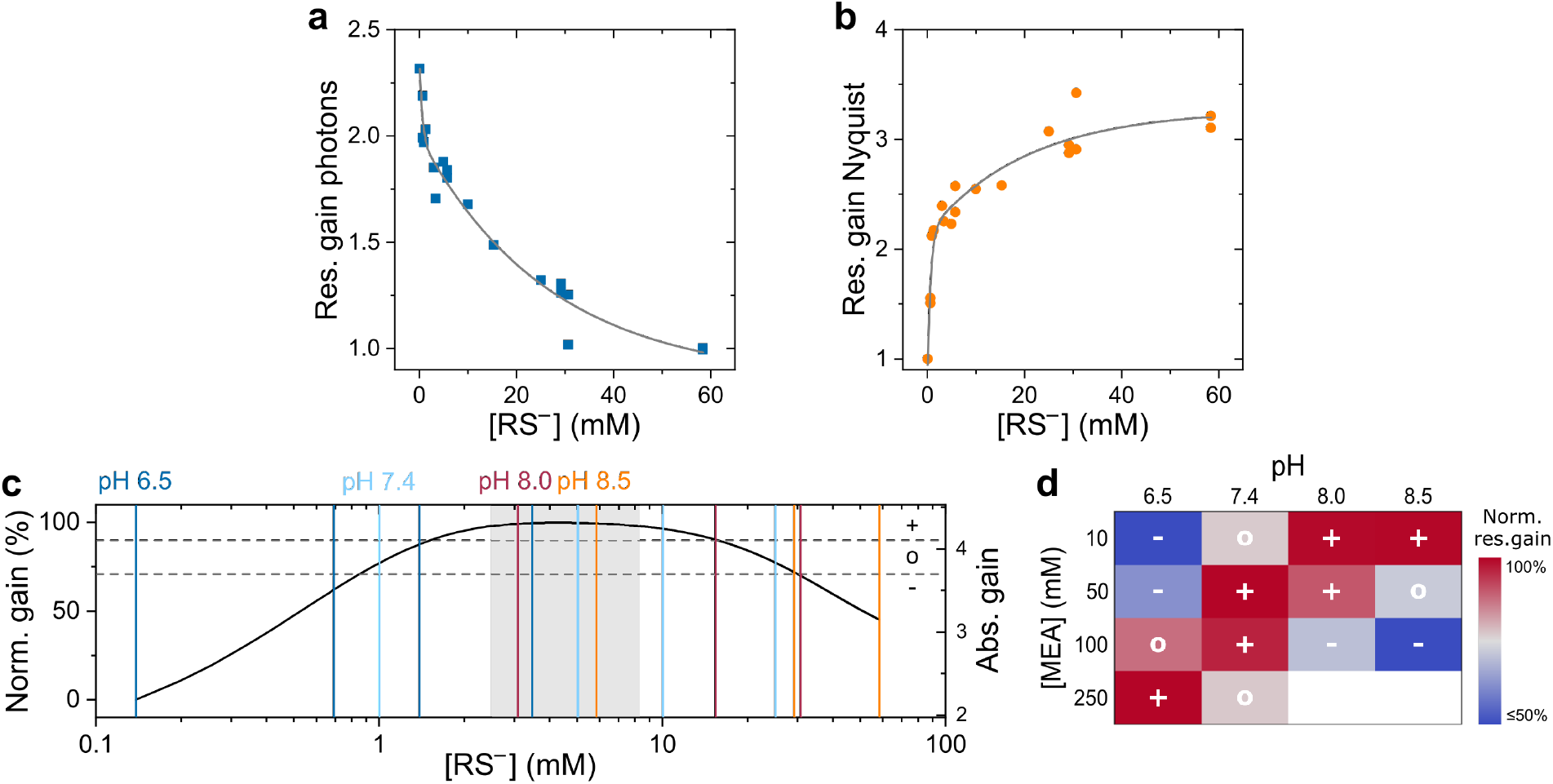
Optimal thiolate concentration. **a, b**) Resolution gain as shown in Fig. 4 but with linear scaling; **a**) resolution as determined by the photon limited localization precision and **b**) Nyquist resolution as determined by the maximum achievable label density, which is defined by the τ_off_*/*τ_on_ ratio. Note the pronounced increase in Nyquist resolution towards 1 mM RS*−*. **c**) Overall gain in resolution as determined by the product of the fit curves as shown in a) and b); left axis shows normalized, right axis absolute gain. Lower horizontal and upper dashed lines indicate 70.7% (-3 dB) and 90% of the total gain, respectively. A working range for the thiolate concentration can be determined between 1.5 and 15.6 mM with 90% total gain, the optimal range can be found between 2.5 and 8.3 mM (98%, light grey area) with 4.3 mM as maximum gain. The concentration bandwidth as determined at -3 dB allows to identify the lowest and highest RS*−* concentration of 0.85 mM and 30.15 mM, respectively. The vertical lines indicate the tested buffer conditions; colour indicates the pH with the MEA concentrations increasing from left to right: 10, 50, 100, (250) mM MEA. Buffer settings below 70.7% are marked *−*, between 70.7% and 90% *∘* and above 90% + in panel d). **d**) Rating of the tested buffer conditions on the basis of the classification as shown in panel c).

**Fig. S7.**
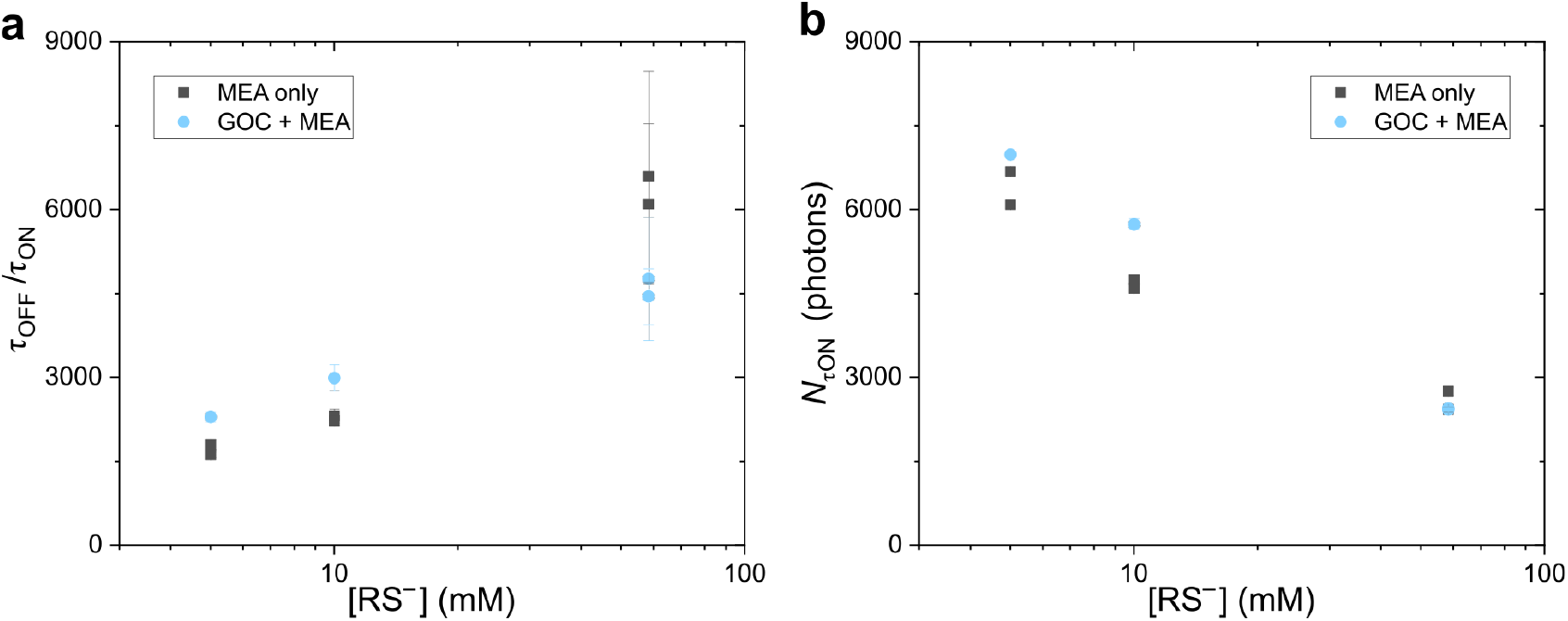
Photoswitching metrics of AF647 with and without enzymatic oxygen scavenger system. **a**) Photon budget *N*τon and **b**) τ_off_*/*τ_on_ ratio as function of thiolate concentration.

**Fig. S8.**
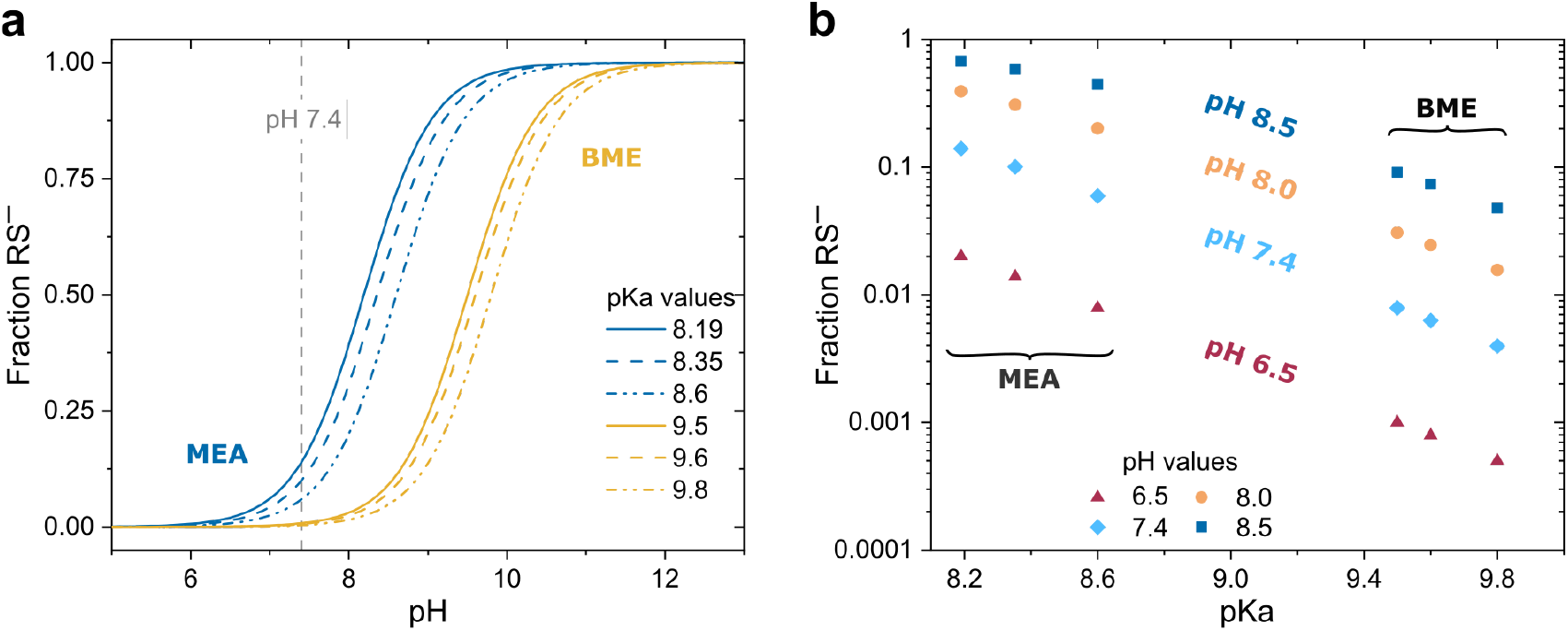
p*K* a dependent fraction of thiolate RS*−*. **a**) Fraction of RS*−* for different p*K* a values as a function of the pH. Curves in blue and orange show typical p*K* a values as published for MEA and BME, respectively. **b**) Fraction of RS*−* for different pH values as a function of the p*K* a. Data points between 8.2 and 8.6 refer to p*K* a values as published for MEA, whereas data points between 9.5 and 9.8 refer to p*K* a values as published for BME.

**Table S1.** pH dependent thiolate concentration for varying MEA concentrations. All concentrations in mM. Thiolate concentrations were determined according to the Henderson-Hasselbalch equation (cf. Eq. (1)) with the p*K* a as determined in Fig. S5.

| [MEA]<br>(mM) | pH |  |  |  |
| --- | --- | --- | --- | --- |
|  | 6.5 | 7.4 | 8.0 | 8.5 |
| 10 | 0.14 | 1.00 | 3.07 | 5.84 |
| 50 | 0.69 | 5.01 | 15.36 | 29.19 |
| 100 | 1.38 | 10.03 | 30.73 | 58.38 |
| 250 | 3.46 | 25.06 | 76.82 | 145.96 |

